# Metagenomic Thermometer

**DOI:** 10.1101/2022.07.14.499854

**Authors:** Masaomi Kurokawa, Koichi Higashi, Keisuke Yoshida, Tomohiko Sato, Shigenori Maruyama, Hiroshi Mori, Ken Kurokawa

**Affiliations:** Genome Evolution Laboratory, National Institute of Genetics, 1111 Yata, Mishima, Shizuoka 411-8540, Japan; Department of Biological Information, Tokyo Institute of Technology, 2-12-1 Ookayama, Meguro-ku, Tokyo 152-8550, Japan; Earth-Life Science Institute, Tokyo Institute of Technology, 2-12-1 Ookayama, Meguro-ku, Tokyo 152-8551, Japan

**Keywords:** metagenome, temperature, optimal growth temperature, hot spring, human gut, community assembly

## Abstract

Various microorganisms exist in environments, and each of which has an optimal growth temperature (OGT). The relationship between genomic information and OGT of each species has long been studied, and one such study revealed that OGT of prokaryotes can be accurately predicted based on the fraction of seven amino acids (IVYWREL) among all encoded amino-acid sequences in its genome. Extending this discovery, the fraction of IVYWREL among all amino acids encoded in the metagenomic DNA in an environment may reflect the environmental temperature to which the microorganisms have adapted. In the present study, we developed a ‘Metagenomic Thermometer’ as a means of predicting environmental temperature based on metagenomic sequences. Analysis of 41 metagenomic samples obtained from hot springs of varying temperature revealed that the Metagenomic Thermometer could indeed predict environmental temperature with high accuracy (*R*^*2*^ = 0.75). Although the taxonomic composition of microbial community was substantially affected by pH, the performance of the Metagenomic Thermometer was refractory to changes in community composition. The Metagenomic Thermometer was also applied to 616 human gut metagenomic samples, yielding a reasonably accurate value (i.e., 36.22 ± 1.98°C) for human body temperature. Our results suggest that environmental temperature can be predicted based on metagenomic sequence data.

## Introduction

The effect of temperature on microbial growth has long been a topic of research^1^. It is generally understood that each microbial species has an optimal growth temperature (OGT), and that growth rate decrease as the temperature deviates from OGT^2,3^. This may be attributable to the temperature sensitivity of the structures of their biomolecules^4^. Therefore, a number of relationships have been discovered between OGT and genomic information. For example, the GC content of microbial tRNAs and rRNAs correlates with OGT^5,6^. Also, compared with mesophiles, thermophiles have a higher proportion of purine bases (A and G) in their mRNAs^7,8^. Moreover, the amino-acid composition of microbial proteins also correlates closely with OGT. For example, overall proportion of charged amino acids is relatively high in thermophiles, which is thought to contribute to salt-bridge formation between negatively charged amino-acid residues and positively charged residues^9,10^. Interestingly, Zeldovich et al. found that the fraction of seven amino acids, namely Ile, Val, Tyr, Trp, Arg, Glu, and Leu (abbreviated *F*_*IVYWREL*_), in a proteome strongly correlates with OGT; and indeed OGT could be accurately predicted by a simple equation with *F*_*IVYWREL*_ as the only variable^11^.

The OGT of a microorganism also reflects the temperature of its natural habitat. Thermophiles, which by definition have an OGT of ≥45°C^12^, are usually found in hot springs. For example, species of the phylum *Chloroflexi*^13–17^ are often the predominant species found in hot springs near 50°C, and *Aquificae*^13,15,16,18,19^ and *Deinococcus-Thermus*^16,18,20^ include the major species in hot springs warmer than 50°C. Therefore, we hypothesized that environmental temperature could be predicted based on the overall *F*_*IVYWREL*_ value for all microorganisms present in a particular habitat.

Here, we report the concept of a ‘Metagenomic Thermometer’, i.e., that the temperature of a sampling site could be predicted using *F*_*IVYWREL*_ obtained for amino acid sequences translated from metagenomic DNA sequences. Hot springs are a suitable environment for confirming the effectiveness of Metagenomic Thermometer because of their wide temperature range. Also, both the temperature and microbiome of a hot spring are generally stable throughout the year^21^. However, the microbial community of a hot spring can be affected by characteristics other than temperature, such as pH^19^ and chemical composition^22,23^. Therefore, we constructed a hot-water canal to create an environment in which conditions other than temperature could be held constant. By applying a Metagenomic Thermometer, the temperature of hot-water canal could be predicted with high accuracy. Further analysis of public metagenomic sequence data acquired for various hot springs revealed that Metagenomic Thermometer is refractory to changes in conditions other than temperature. Finally, we tested whether Metagenomic Thermometer could be applied to the human gut metagenomic data. Overall, the results demonstrate that the temperature of an environment can be predicted based on DNA information derived from the environment and provide important insights for understanding the contribution of temperature to the composition of microbial communities.

## Materials & Methods

### Construction of a hot-water canal

A hot-water canal was constructed in February 2014 to enable experimentation at different water temperatures. Water for the canal was drawn from Kin-yu hot spring, which is located in the Kirishima area in central Kagoshima prefecture in Japan. The Kirishima area is located on the volcanic front where the Philippine Sea plate sinks into the Eurasian plate^24^, and approximately 20 volcanoes exist in the area of approximately 20 km × 30 km^25^. Thus, this region has numerous hot springs that range in temperature from approximately 40°C to 97°C and in pH from approximately 2 to 9^22,26,27^. Kin-yu hot spring has a temperature range of 70–85°C with near-neutral pH^22,27^.

The canal length was ∼20 m and was created beside the Kin-yu source using U-shaped gutters (Fig. 1). Water was drawn from the source through a pipe, and the downstream outlet was naturally open. A net was placed on the canal to prevent clogging by leaves, yet soil and microbes could enter freely.

**Fig. 1.**
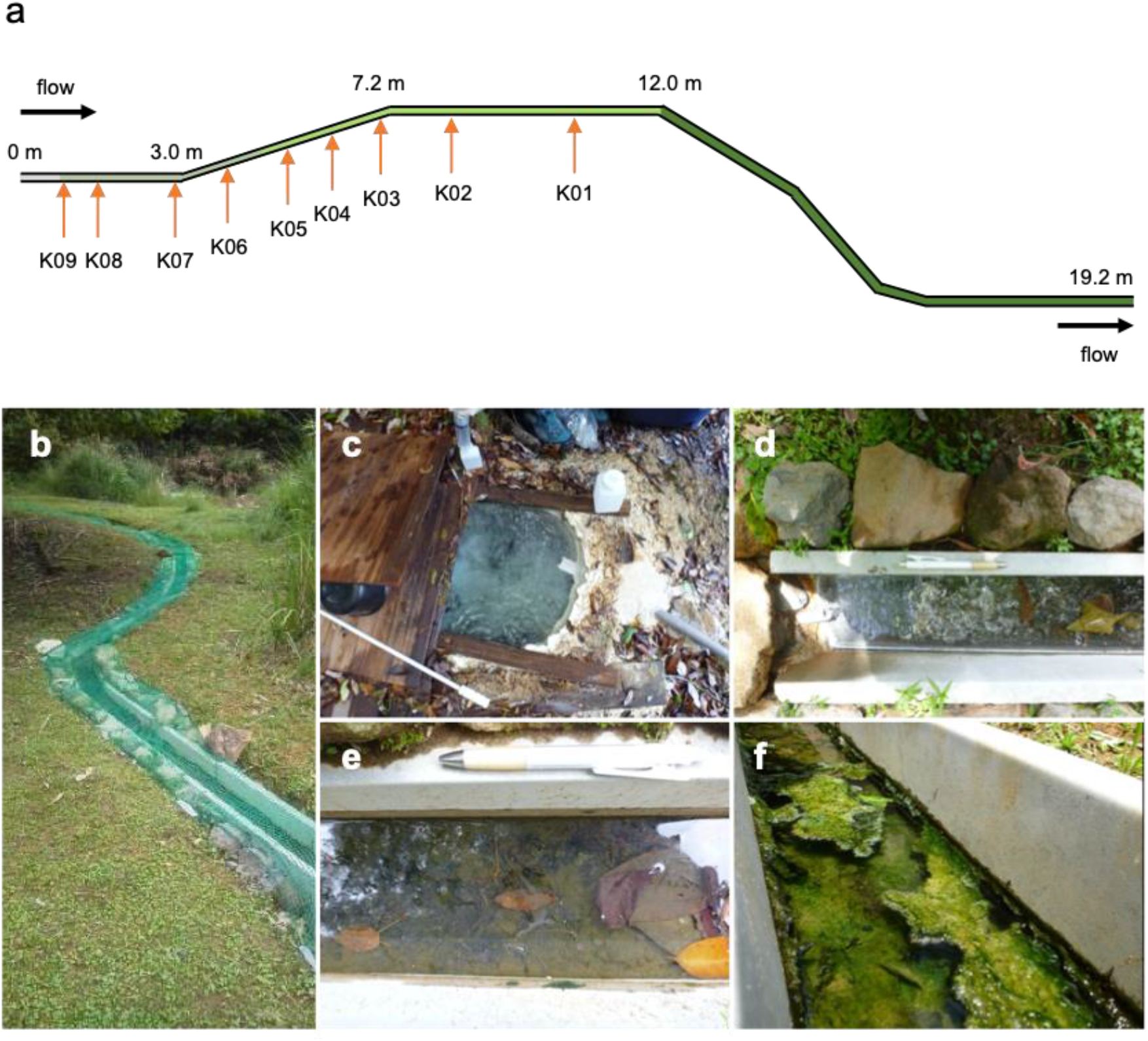
Schematic diagram of a hot-water canal. **a** The elongated line represents the hot-water canal. Water flow was from left to right, and the distance from the starting point of the flow is shown above the line. Sediment color was yellow or gray up to ∼1 m, dark gray or dark green up to ∼4 m, light green up to ∼12 m, and light green downstream from there. Orange arrows indicate the sampling points. **b** Photograph of the canal viewed from a downstream point. **c** The source of the Kin-yu. The pipe leads to the canal. **d** Upstream of canal connected to the source by a pipe. **e** Sediment with a light-green biomat. **f** Sediment with a thick, light-green biomat.

### Sample collection

All samples from the canal were collected in February 2016. Surface-sediment samples (∼15 g) from the canal were placed into 50-mL conical tubes using sterilized steel spatulas. To avoid contamination between downstream and upstream sediment, sampling was carried out from downstream to upstream. Also, different spatulas were used for each sampling. Collected samples were immediately placed in a cooler and transported to the laboratory, where they were stored at −20°C. Immediately before sampling, the pH and temperature of the canal water were measured using a portable conductivity/pH meter (WM-32EP, DKK-TOA Co.), and oxidation-reduction potential and dissolved oxygen of the water were measured using a portable dissolved oxygen/pH meter (DM-32P, DKK-TOA Co.) at a site just above each sampling point. Sampling was carried out in three replicate at each point. One was for shotgun sequencing, another was for amplicon sequence, and the other was as a spare.

### DNA extraction

Frozen sediment samples were thawed at room temperature, and DNA was extracted from the samples using the PowerSoil DNA Isolation kit (MO BIO; version 01152013) with a Micro SmashMS-100R (TOMY) for bead homogenization (3,000 rpm, 30 s). Each sample of extracted DNA was in a volume of 50 µL. DNA concentration was initially approximated using a Qubit 2.0 Fluorometer (Thermo Fisher Scientific), with subsequent accurate measurement using a Model 2100 Bioanalyzer (Agilent Technologies).

### PCR amplification of the 16S rRNA gene

From the extracted DNA, the V3-V4 region of the 16S rRNA gene was amplified using universal primers 342F and 806R^28^. PCR mixtures contained 21 μL sterilized distilled water, 3 μL template DNA solution, 25 μL Premix Taq Hot Start Version (TaKaRa), 0.5 μL forward primer, and 0.5 μL reverse primer. The thermal cycler protocol was 95°C for 2 min, followed by 30 cycles of 98°C for 10 s, 50°C for 30 s, and 72°C for 40 s, with final extension at 72°C for 10 min. PCR products were separated by electrophoresis (1.5% agarose gel) and purified using the Wizard SV Gel and PCR Clean-Up System (Promega). Purified DNA was quantified using the Model 2100 Bioanalyzer with a High Sensitivity DNA kit (Agilent Technologies; version 01.02).

### Sample preparation for shotgun metagenomic sequencing

Extracted DNA samples were processed using the Nextera DNA Library Preparation kit (Illumina) per Nextera DNA Sample Preparation Guide #15027987 and purified using the DNA Clean & Concentrator™ (ZYMO RESEARCH). Finally, libraries were subjected to quantitative PCR using the KAPA Library Quantification kit (NIPPON Genetics Co., Ltd.) per manufacturer’s protocol for Illumina (2013.9.6 rev6).

### DNA sequencing

Paired-end DNA sequencing was performed by Earth-Life Science Institute (Tokyo Institute of Technology) using MiSeq (Illumina) per the MiSeq System Quick Reference Guide For MiSeq Software 2.3. In the 16S rRNA amplicon sequence, PhiX was spiked into the DNA samples at 50% relative concentration for quality control. Metagenomic sequencing reads were deposited in the DNA Data Bank of Japan with accession number DRA012733.

### Acquisition of public sequence data

A total of 33 hot-spring metagenomic samples, 616 human gut metagenomic samples, and 586 ocean metagenomic samples were used for comparative metagenomic analyses. All public metagenomic datasets were downloaded from the NCBI Sequence Read Archive database using SRA Toolkit.

### Quality control of sequence reads

Adapter trimming and removal of low-quality reads were performed using Fastp v0.20.0^29^. The lower limit of read length was set to 50 bp, and the allowable limit of the number of ambiguous nucleotides (N) per read was set to 1.

### Acquisition of published information for OGT

The OGT for each bacterial species based on actual measurements was obtained according to the online dataset https://doi.org/10.5281/zenodo.1175608. This dataset was summarized by Engqvist and published in 2018^30^. This dataset was constructed by integrating information from six major databases and contains OGT data for 21,498 nonredundant microbes.

### Prediction of OGT

The OGT for each bacterial species was predicted based on the method reported by Zeldovich et al^11^. Amino-acid sequence fasta files for coding regions of focal bacterial species were downloaded from the NCBI Genome database. *F*_*IVYWREL*_ was calculated for all coding sequences of focal species, and then OGT for each species was predicted by applying the following formula (Eq. 1).

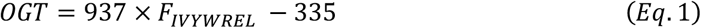

*Calculation of the metagenomic predicted temperature (MPT) and implementation of web application* Quality-filtered metagenomic DNA fastq files were converted to DNA fasta files using SeqKit v0.16.1. Using Prodigal v2.6.3, open reading frames were identified from DNA fasta files, which yielded the corresponding amino-acid fasta files. *F*_*IVYWREL*_ values were calculated from the amino-acid fasta files, and Eq. 1 was applied to calculate MPT instead of calculating OGT. Hereafter, we call this temperature prediction method the Metagenomic Thermometer, for which a web application (written in Python) is freely available at http://metathermo.jp/. Users can upload a fasta or fastq file for shotgun sequencing data and receive a corresponding MPT value.

### Taxonomic profiling of microbial communities

The taxonomic composition of each metagenomicsample was determined using VITCOMIC2^31,32^ based on metagenomic DNA data in the fasta files (only forward read files). VITCOMIC2 identifies 16S rRNA gene sequences from input data and returns results for both genus- and phylum-level taxonomic composition. The abundance of each taxon in each sample was converted to a relative value for each sample and used for subsequent analysis.

Complete linkage–based hierarchical clustering was performed against the genus-level taxonomic compositions. The distance between each microbial community was defined as the Bray-Curtis dissimilarity calculated using the R^33^ package vegan^34^. The R package program dendextend^35^ was used to visualize dendrograms.

### LEA mapping

To analyze the characteristics of the microbial community structure, metagenomic DNA sequence data in fasta files for 616 human gut metagenomic samples were analyzed by VITCOMIC2. The resulting files provided by VITCOMIC2 were uploaded to the LEA web application (http://leamicrobe.jp)^36^, and metadata for each sample were manually annotated and visualized.

## Results

### Physicochemical properties of the hot-water canal samples

Table 1 lists the physicochemical conditions at nine sampling points (K-01 to K-09, downstream to upstream) in the experimental hot-water canal (Fig. 1a). As expected, the temperature dropped from upstream (60.0°C) to downstream (29.6°C). The pH range was 7.62–8.31 (weakly alkaline) over all sampling points. Dissolved oxygen increased with decreasing temperature (Fig. S1a), perhaps because oxygen solubility is higher at lower temperature. The oxidation-reduction potential correlated negatively with temperature and positively with dissolved oxygen (Fig. S1b, c).

**Table 1.** Physicochemical properties of samples.

| Sampling point | Temperature (°C) | pH | ORP (mV) | DO (mg/L) |
| --- | --- | --- | --- | --- |
| K-01 | 29.6 | 8.00 | −19 | 0.97 |
| K-02 | 34.7 | 7.99 | −19 | 0.79 |
| K-03 | 36.8 | 8.01 | −54 | 0.87 |
| K-04 | 40.0 | 8.31 | −86 | 0.84 |
| K-05 | 43.1 | 8.24 | −105 | 0.58 |
| K-06 | 44.6 | 8.28 | −104 | 0.56 |
| K-07 | 50.2 | 8.28 | −134 | 0.57 |
| K-08 | 54.0 | 7.91 | −180 | 0.34 |
| K-09 | 60.0 | 7.62 | −272 | 0.49 |

DO: dissolved oxygen
ORP: oxidation-reduction potential

### Microbial community composition

A total of 48,150,715 high-quality shotgun metagenomic reads with average length 181 bp were obtained from eight of the nine hot-water canal samples (for K-04, the DNA concentration was insufficient for shotgun metagenomic sequencing). The phylum-level taxonomic composition for each sample was examined using VITCOMIC2 (Fig. 2a). Each sample had 21–31 bacterial phyla, and 34 different phyla were represented in total. *Chloroflexi* and *Proteobacteria* were the predominant phyla in the microbiome among all samples, representing ∼34% and 33% of the population on average, respectively, followed by two other phyla, *Bacteroidetes* (4.2%) and *Aquificae* (3.8%). Note that Proteobacteria was classified at the class level (e.g., *Alphaproteobacteria, Betaproteobacteria*, etc.) according to the VITCOMIC2 results. Previously, a comprehensive comparison of an entire bacterial community, as derived from metagenomic data for 30 hot springs in Yellowstone National Park, revealed *Chloroflexi* as the predominant phylum in hot springs with pH above 5 and temperature below 70°C, consistent with our result^13^. That study also found that *Aquificae* predominated in flowing channels^13^. Also, *Proteobacteria* have frequently been found in various hot springs having a wide range of temperatures^37–40^. Although *Bacteroidetes* is abundant in the human gut^41^, they often exist in hot springs of moderate temperature^39,40,42^. The results of the 16S rRNA gene amplicon sequencing (performed on samples from the same hot-water canal site for the shotgun analysis) are shown in Fig. S2. Notably, K-04 was not an abnormal sample among the other eight samples.

**Fig. 2.**
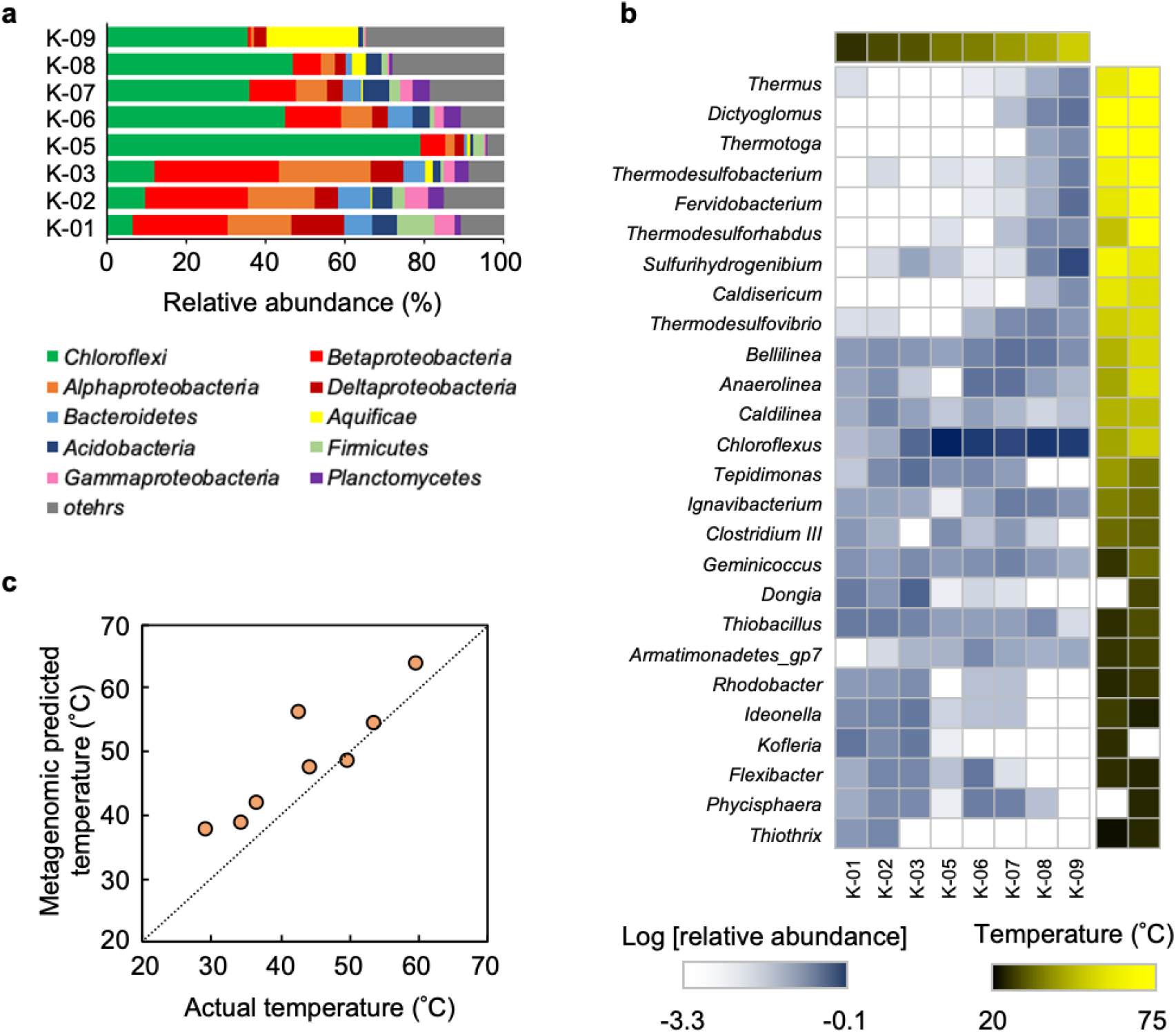
Analysis of the microbial community of the experimental hot-water canal. **a** Relative abundance of microbial phyla/class in each sample. Colors correspond to the phyla or class (for *Proteobacteria*) as shown in the legend. Labels on the left represent the sample points of the canal indicated in Fig. 1a. **b** Heatmap of major genera abundance. Genera with relative abundance of more than 2% in any sample are shown. Each column represents a single sampling point, and the color of the above cells indicates the temperature of the samples correspond to the scale. The two columns of colored squares shown on the right indicate the median OGT for the species included in the genus; the left column indicates the optimum temperature obtained based on the datasets, and the right column indicates the temperature predicted based on *F*_*IVYWREL*_. **c** Plot of MPT versus actual temperature in eight of the nine canal sampling points. The dotted line represents the perfect match between the predicted and actual temperatures.

To better understand the effect of temperature on microbial community composition, we investigated the relationship between the temperature of sampling point and OGT of the predominant genus. For genera representing >2% in at least one sample, the median OGT of the species belonging to that genus was investigated. OGT was obtained from the public dataset and also predicted using *F*_*IVYWREL*_ values from genetic information (for details, see Materials & Methods). The results revealed that genera having a relatively higher OGT tended to be more abundant in higher-temperature samples, and vice versa (Fig. 2b). Therefore, although OGT is defined by cultures in laboratory, it also seems to reflect relative prosperity of a bacterial species in nature. Furthermore, certain genera were found in abundance even though their OGT differed substantially from the temperature of environment in which they were found.

### Metagenomic Thermometer can predict temperature of sampling sites

To verify whether environmental temperature can be predicted by the Metagenomic Thermometer, we calculated MPT as described in Materials & Methods. The results revealed that temperature of hot-water canal samples could be predicted based on metagenomic sequence information, with an accuracy of 6.00°C root-mean-square error (*RMSE*) and a coefficient of determination (*R*^*2*^) of 0.61 (Fig. 2c, orange circles). One exception was sample K-05, for which the predicted temperature was significantly higher than the actual temperature. This may be attributable to the large proportion of *Chloroflexi* in K-05 compared with other samples (Fig. 2a). The average predicted OGT of species in *Chloroflexus*, the predominant genus of *Chloroflexi* in K-05, was ∼60.2°C (Fig. 2c). The extraordinary abundance of *Chloroflexi* in K-05 is probably attributable to sample-collection bias, such as contamination with biomat debris. Analysis of the 16S rRNA amplicon sequences of K-05 sample, which is independently collected replicate sample from the same sampling point, showed that the relative abundance of *Chloroflexi* in K-05 was not very high (Fig. S2). For reference, the prediction accuracy of sample temperature is improved using seven samples except for K-05 (R^2^ = 0.83).

### Metagenomic Thermometer can be applied to hot springs of diverse characteristics

To assess the ability of the Metagenomic Thermometer to predict water temperature in hot springs of diverse chemical and geological characteristics, we utilized public metagenomic shotgun sequencing data for a variety of hot springs. As of May 2021, the NCBI Sequence Read Archive database contained shotgun metagenomic sequences for 39 hot-springs with temperature data. We removed samples having a fastq file size of <10 MB or if contamination of the biological sample was suspected, which left 33 samples for analysis. A diversity of hot springs was assessed, with sampling points from six countries and a pH range of 1.9–9.8. The Metagenomic Thermometer could reasonably predict the temperature of the hot springs based on the metagenomic sequence data (*RMSE* = 10.87, *R*^*2*^ = 0.75) (Fig. 3a).

**Fig. 3.**
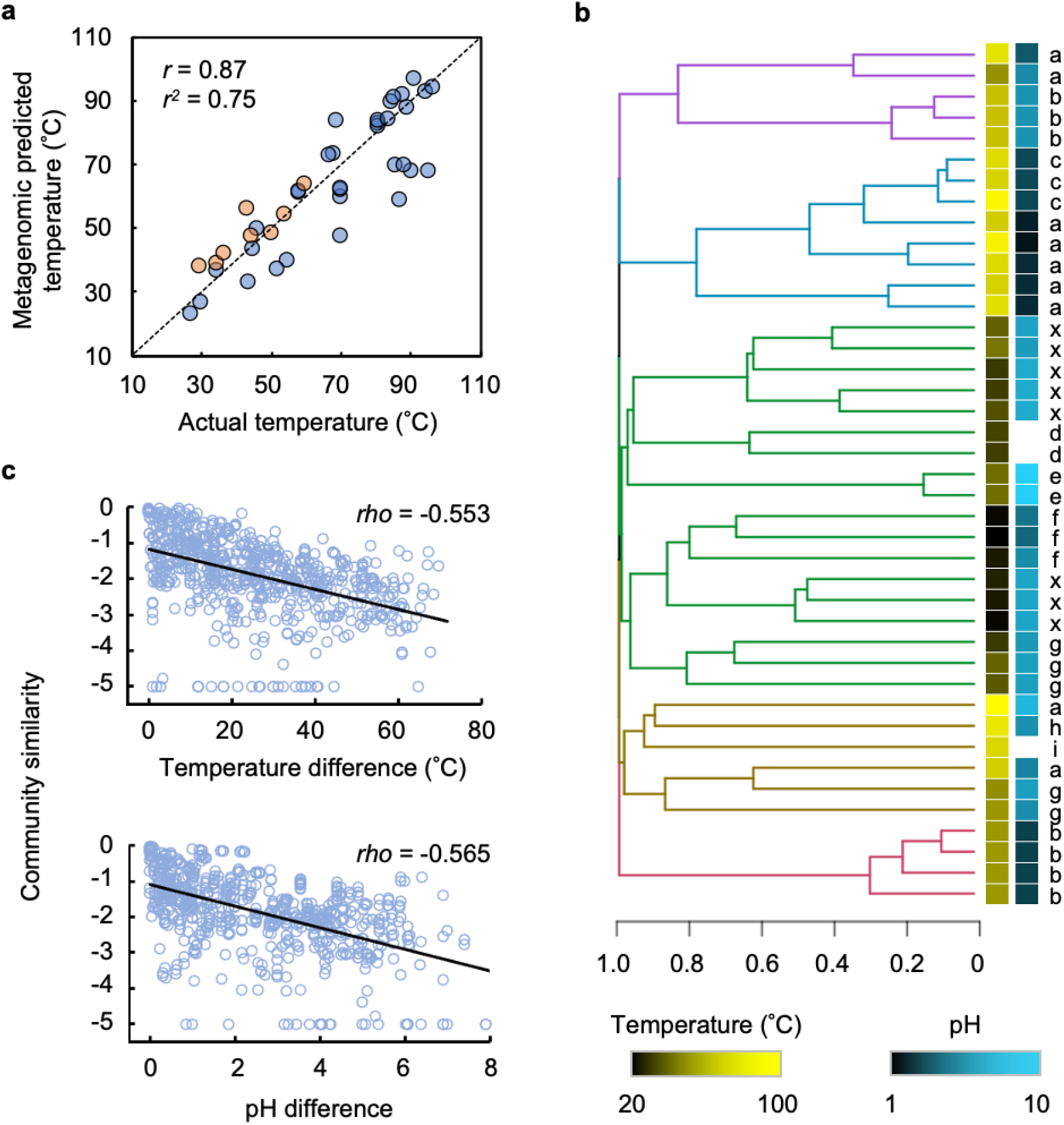
Temperature prediction of a variety of hot springs. **a** Plot of MPT values of 33 public hot-springs (blue) or eight sampling points along the experimental hot-water canal (orange, same as Fig. 1c) versus actual temperature. The results were significantly correlated (*p* = 7.63 × 10^−14^). The dotted line represents the perfect match between the actual and predicted temperatures. **b** Dendrogram showing the clustering of hot-springs samples having a similar genus-level community structure. The yellow and blue heatmaps show the temperature and pH, respectively, of each hot spring. No pH information could be obtained from the three data. Each ‘x’ corresponds to data obtained in this study. Branches are colored for each six cluster separated by the k-mean method. Scale bar representing the relative taxonomic distance. **c** Pairwise community similarity decreases as the difference in temperature or pH between samples increases (*p* < 2.2 × 10^− 16^). ‘Community similarity’ was calculated as 1 minus the calculated Bray-Curtis dissimilarity.

Based on the success of the Metagenomic Thermometer, the question arose whether the microbial community structure could be characterized based on environmental temperature. This issue was addressed via a clustering analysis of the community taxonomic composition (Fig. 3b). Inspection of the taxonomic dendrogram revealed that the microbes sampled from relatively low-temperature springs were grouped in the same cluster (green branches in Fig. 3b), whereas those taken from relatively high-temperature springs were classified into four clusters. The high-temperature samples could be further divided into low- and high-pH groups from relatively low-temperature environments (pink and purple branches) or from very high-temperature environments (blue and ocher branches). These results revealed that pH is also an important determinant of community structure.

To examine the relative contribution of temperature and pH to community structure, the relationship between the difference in temperature or pH and the similarity of the microbial community structure for pairs of samples was analyzed (Fig. 3c). The results indicated that the magnitude of the temperature or pH difference correlated with the relative difference in community structure, suggesting that both temperature and pH can affect community structure. We speculated that samples taken from sites proximal to one another would have similar temperature and pH values and hence a similar community structure, and thus another analysis was performed only on pairs of samples from different countries to eliminate any geographical bias; however, the results were actually similar (Fig. S3). These results suggested that Metagenomic Thermometer can predict the temperature of the sampling site even though the community structure is greatly affected by factors other than temperature, such as pH.

### Application of the Metagenomic Thermometer for prediction of the temperature of the human gut

To confirm whether the Metagenomic Thermometer could be applied to environments other than hot springs, MPT was calculated based on metagenomic data for the human gut. The metagenomic data was obtained from the DNA Data Bank of Japan (DRA PRJDB4176) that consisted of 616 individuals including healthy subjects and patients with colorectal cancer^43^. This dataset was suitable for our study owing to the large number of samples, and indeed metadata were available for individuals. The MPT value calculated based on all samples revealed an average value of 36.22 ± 1.98°C (standard deviation). Given that the average human body-temperature range is 36–37°C^44,45^, the Metagenomic Thermometer could predict body temperature with high accuracy. Stratification of the data by gender revealed that the predicted temperature tended to be higher for females (36.45 ± 1.66°C) than males (36.02 ± 2.16°C) (Fig. 4a). Large-scale research on human body temperature has also shown that female body temperature tends to be higher than that of males^44,46^. Body temperature also has been reported to correlate with age^44,45^ and body mass index^44^, but our results indicated no correlation between predicted temperature and these two parameters (Fig. S4). Notably, our data revealed no difference in the ability of the Metagenomic Thermometer to predict body temperature in colorectal cancer patients versus healthy subjects (Fig. 4b).

**Fig, 4.**
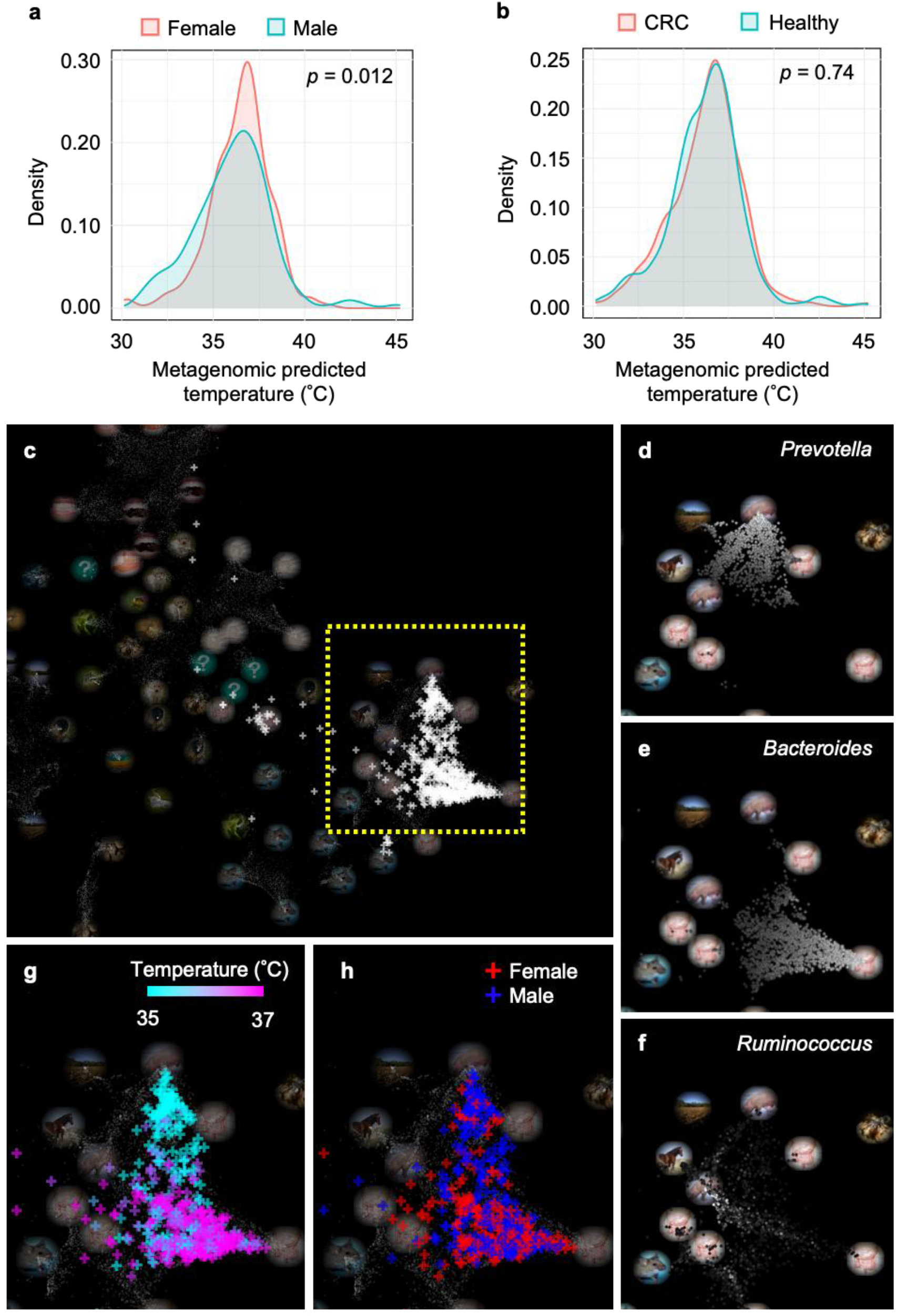
Temperature prediction based on the human gut microbiome. **a, b** Probability density distribution of MPT calculated based on the gut microbiome is shown separately according to gender (**a**) and whether an individual had colorectal cancer (CRC) (**b**). **c** LEA map with all 616 samples (white plus signs). The various circular photographs represent environmental topics, that corresponds to microbial communities that represent the latent environment. Samples or environmental topics that are relatively close in the LEA space have similar community structures. The area enclosed by the yellow dotted rectangle is enlarged in panels **d–h. d–f** The myriad points represent metagenomic samples stored in MicrobeDB, and the higher the proportion of *Prevotella* (**d**), *Bacteroides* (**e**), and *Ruminococcus* (**f**) in each sample, the whiter is the sample color. **g, h** Samples used in this study are color coded according to predicted temperature (**g**) or gender (**h**).

To assess how a difference in microbiome could affect the MPT, all samples were mapped to the LEA map. LEA is a web application that visually expresses microbiome similarities between samples and the strength of their relevance to environmental topics^36^. Most of the samples were placed around the environmental topics representing the gut (Fig. 4c). The human gut microbiome can be classified into three enterotypes based on whether the predominant genus is *Prevotella, Bacteroides* or *Ruminococcus*^47^. Visualizing the abundance of these three genera in the public data contained in the LEA map revealed that the three enterotypes could be clustered in different areas on the LEA map (Fig. 4d–f). Visualization of MPT of samples by color scale revealed that samples with a relatively low MPT clustered with the *Prevotella*-dominated enterotype, and samples with a relatively high MPT clustered with the *Bacteroides*-dominated enterotype (Fig. 4g). The low predicted temperature of the *Prevotella*-dominated enterotype was consistent with the lower predicted OGT of *Prevotella* (26.48 ± 3.27°C) compared with that of *Bacteroides* (34.46 ± 2.42°C). Finally, we investigated whether gender affected the distribution of samples on the LEA map. The results revealed a larger number of males than females in the *Prevotella*-dominated enterotype area, which is consistent with the result that the *Prevotella*-dominated cluster had a larger proportion of males in the study that produced this dataset^43^.

## Discussion

Our results demonstrate that the Metagenomic Thermometer can predict the temperature of an environment based on the *F*_*IVYWREL*_ value for coding-sequences of a metagenomic sequence dataset. The results obtained for numerous hot-springs samples show that environmental temperature can be predicted with high accuracy. Still, the Metagenomic Thermometer is not applicable to some environments. For example, we calculated MPT using ocean metagenomic data, but the MPT values did not match well with the actual measured environmental temperatures (*RMSE* = 12.1, *R*^*2*^ = −1.51) (Fig. S6). It might be related to the continuous flux of microorganisms into and out of environments of different temperature. The surface temperature of the ocean changes greatly with latitude and depth^48,49^. The ocean circulates horizontally and vertically; consequently, microbes can drift with the circulation^50^. The ocean microbial community may be affected not only by currents but also by the influx of microbes from environments other than ocean. A large-scale exploration of ocean metagenomes revealed that the genetic composition of the ocean metagenomes is similar to that of the human gut metagenomes, despite the obvious differences in the physicochemical properties of these two environments^51^. It is possible that the influx of bacteria from human and animal feces is a factor in raising the MPT for the ocean metagenomes.

The accuracy of the Metagenomic Thermometer is a matter to be considered. In this study, MPT of the human gut was close to the normal human body temperature. In addition, MPT was consistent with the well-known fact that body temperature of human females is higher than that of males. Although, it is not clear how accurately the body temperature of an individual can be predicted because actual body temperature data is not available for this dataset. Special instruments such as electronic capsules are needed to measure the actual temperature in the intestine^52^. As such, the Metagenomic Thermometer may represent a convenient method for estimating the temperature in the human intestine. It is also very interesting that the male samples were placed around the *Prevotella*-dominated enterotype on the LEA map. The predominance of *Prevotella* in the male gut has been reported in several studies^53–55^. Although there is a study suggesting a sex hormone relationship to *Prevotella* abundance^56^, exact cause of *prevotella* predominance in the male gut remains unclear^53–55^. Our results indicate that body temperature may be a factor in causing *Prevotella* predominance in the male gut.

It is worth pondering why temperature of a sampling site can be predicted by a very simple indicator, i.e., fraction of amino acids fraction. The simplest answer is that the fraction of amino acids corresponds to the OGT, and microbes whose OGT is close to the environmental temperature have a relatively high rate of proliferation. However, the temperature predicted based on an entire metagenome is more accurate compared with the OGT predicted for the most predominant taxa in the environment. For example, the predicted OGT of *Prevotella* (26.48 ± 3.27°C) was significantly lower than human body temperature. In addition, microbial taxa with a wide range of OGT coexist at each sampling point in the canal, even if we focus only on the genera for which abundance in each sample was >2% (Fig. 2b). It is also interesting that the Metagenomic Thermometer was refractory to changes in taxonomic composition (Fig. 3b). The reliability of temperature prediction even with taxon variation can be partly explained by the lottery model. This model is based on the idea that resident species in an environment are species that arrived by chance, and any species could be selected from a pool of species that are able to grow in that environment^57,58^. Extrapolated to the present study, resident species in an environment are randomly selected from a potential pool of species having an fraction of amino-acid suitable for the environmental temperature; however, it is not completely random but is also defined by selective pressures other than temperature, such as pH. Importantly, the selective pressure of the fraction of amino acids imposed by temperature appears to be little affected by pH. Because temperature cannot be regulated across the bacterial cell membrane, all intracellular proteins are affected by the external temperature. On the other hand, acidophiles cope with acidity by keeping intracellular pH close to neutral^59^, so it is not important that any particular intracellular protein acquires acid resistance. The understanding of community assembly mechanisms has long been debated, yet a unified understanding has not been reached^60^. One study revealed that niche-specific deterministic community assembly processes have a greater effect on gene function than on overall taxonomy^61^. Our study underscores the importance of seeing microbial communities not only as a collection of species but also as a collection of more fundamental biomolecules such as nucleotides or amino acids.

## Acknowledgements

We greatly appreciate the entire staff of Nonoyu-Onsen (Energy Earth Corp.) for allowing us to build the hot-water canal and providing water sample from Kin-yu. This work was supported by a Grant-in-Aid for Scientific Research on Innovative Areas “Hadean Bioscience” (No. 26106001), JSPS KAKENHI Grant Number 22H04925 (PAGS), and the National Bioscience Database Center (NBDC) of the Japan Science and Technology Agency (JST). Computations were partially performed on the NIG supercomputer at ROIS National Institute of Genetics.

## Author Contributions

M.K., K.H., H.M., and K.K. were involved in study design and data interpretation. M.K., K.H., and H.M. were involved in the data analysis. K.Y., T.S., S.M., H.M., and K.K. collected the samples. K.Y. performed biological experiments. M.K., K.H., H.M., and K.K. wrote the manuscript.

## Competing Interests statement

The authors declare no competing interests.

**Fig. S1.**
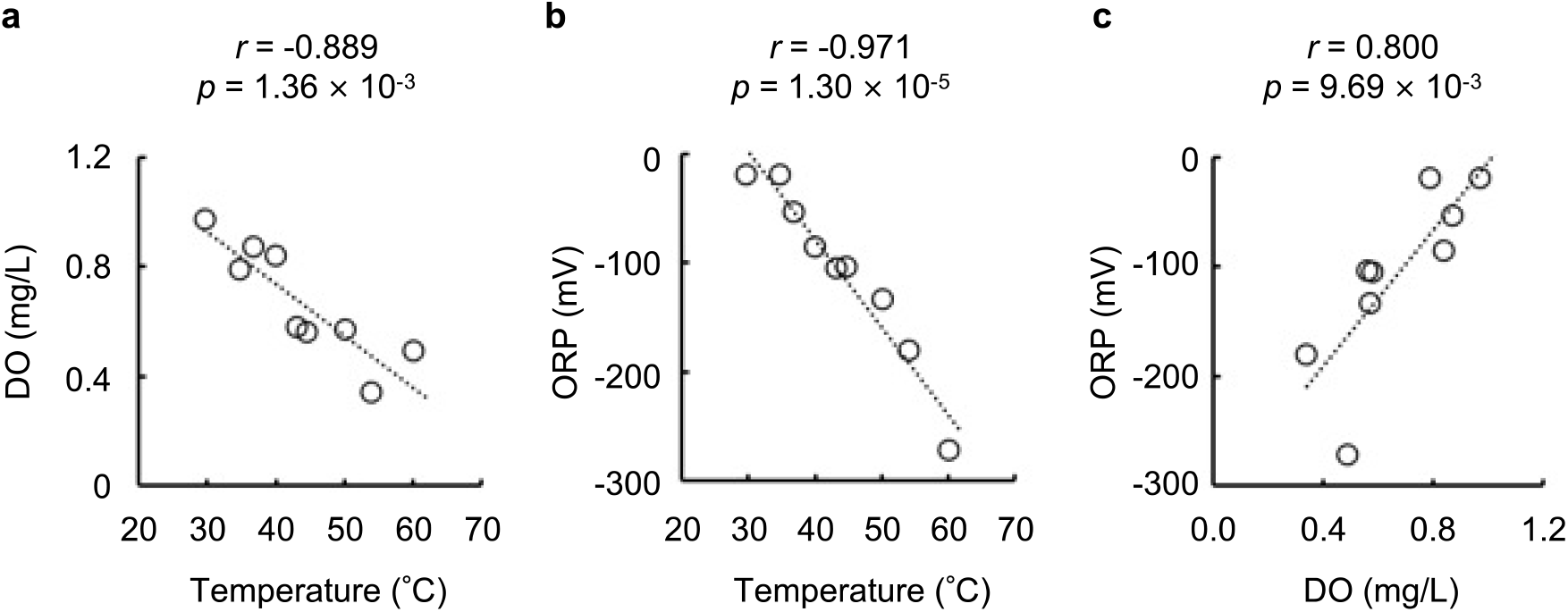
Correlation of physicochemical properties in hot-water canal samples. **a** Temperature and DO (dissolved oxygen) are negatively correlated. **b** Temperature and ORP (oxidation-reduction potential) are negatively correlated. **c** DO and ORP are positively correlated. The Pearson’s correlation coefficient and *p*-value are shown above each graph.

**Fig. S2.**
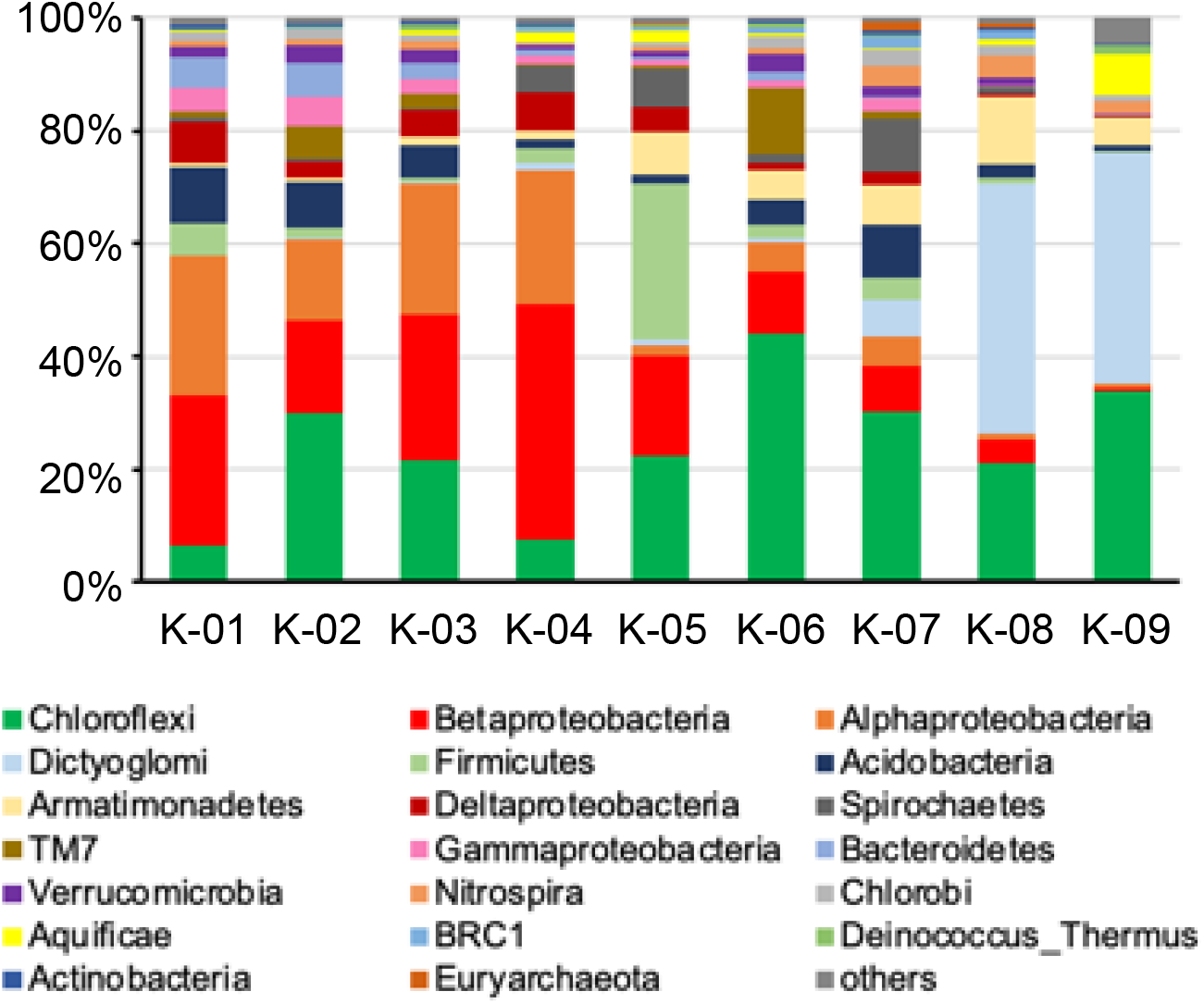
Community structure examined by 16S rRNA gene amplicon sequencing. The 20 most abundant phyla or class (for *Proteobacteria*) in all samples and the remaining phyla correspond to the colors of the legend. Labels at the bottom represent the samples.

**Fig. S3.**
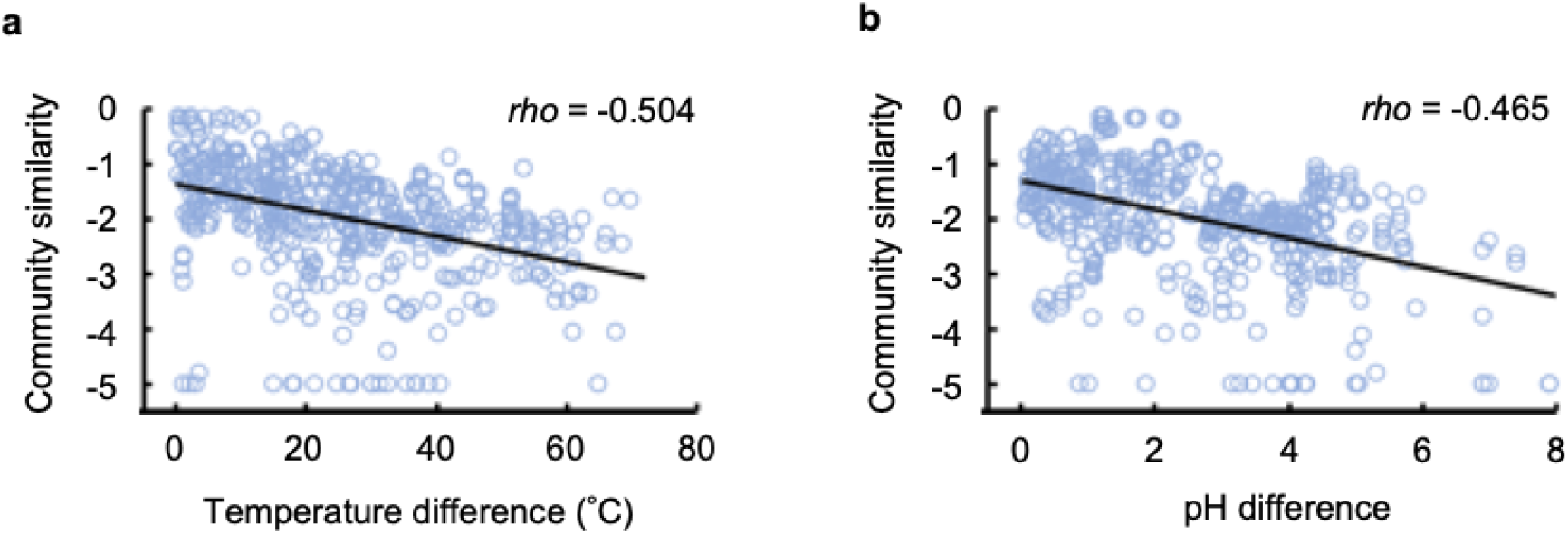
Distance decay graph of pairwise community only different country. Pairwise community similarity decreases according to difference of temperature (a) or pH (b) between samples increases (*p* < 2.2 × 10^−16^). ‘Community similarity’ was calculated as 1 minus the calculated Bray-Curtis dissimilarity.

**Fig. S4.**
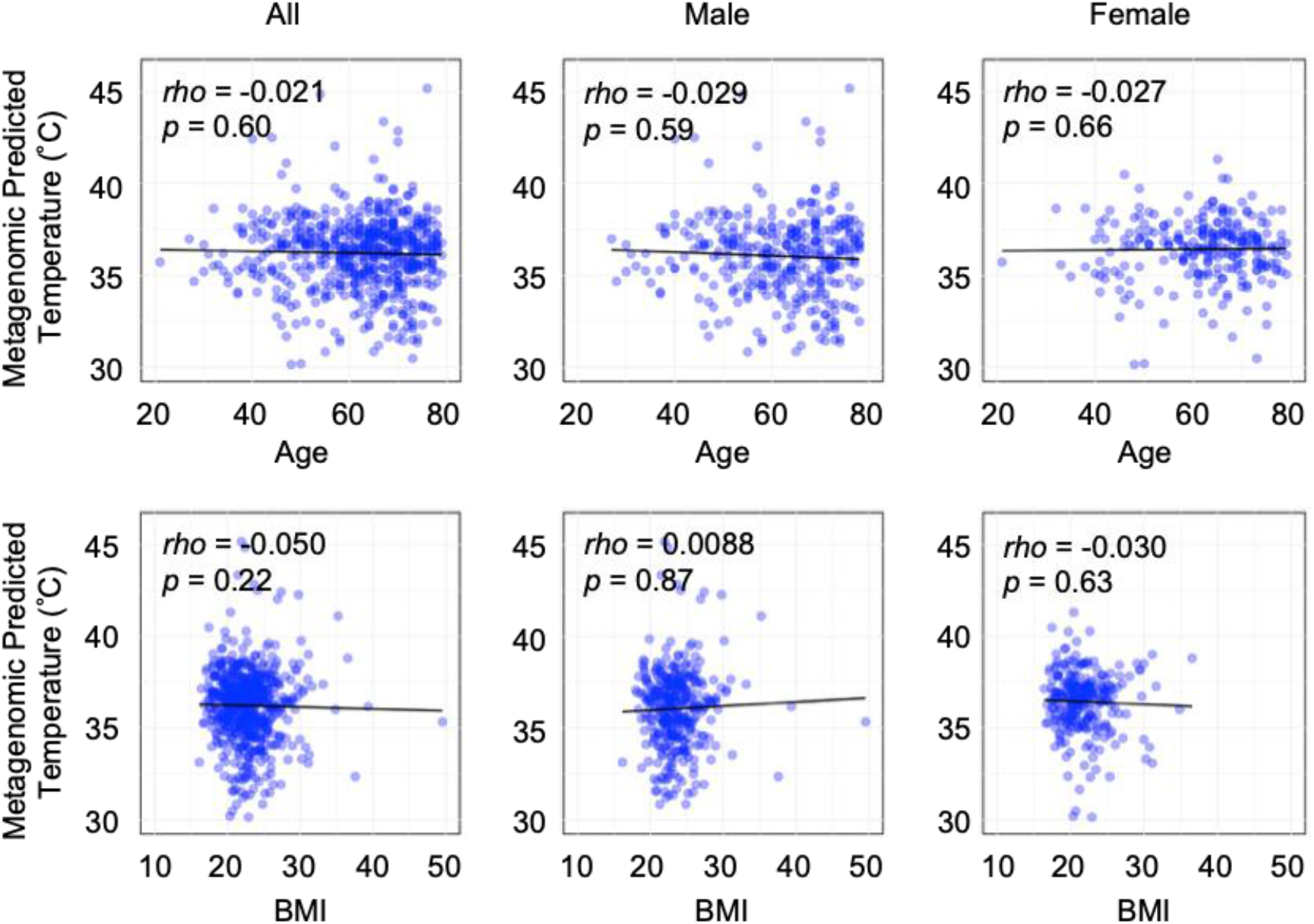
Correlation analysis between predicted temperature and individual characteristics. Correlation between MPT and age (upper) and body mass index (BMI; lower). The left-most graphs present the data for all samples, whereas the middle and right graphs present data for samples pertaining to males and females, respectively. Each black line indicates the linear regression line. Spearman’s correlation coefficients and *p*-values are shown in each graph.

**Fig. S5.**
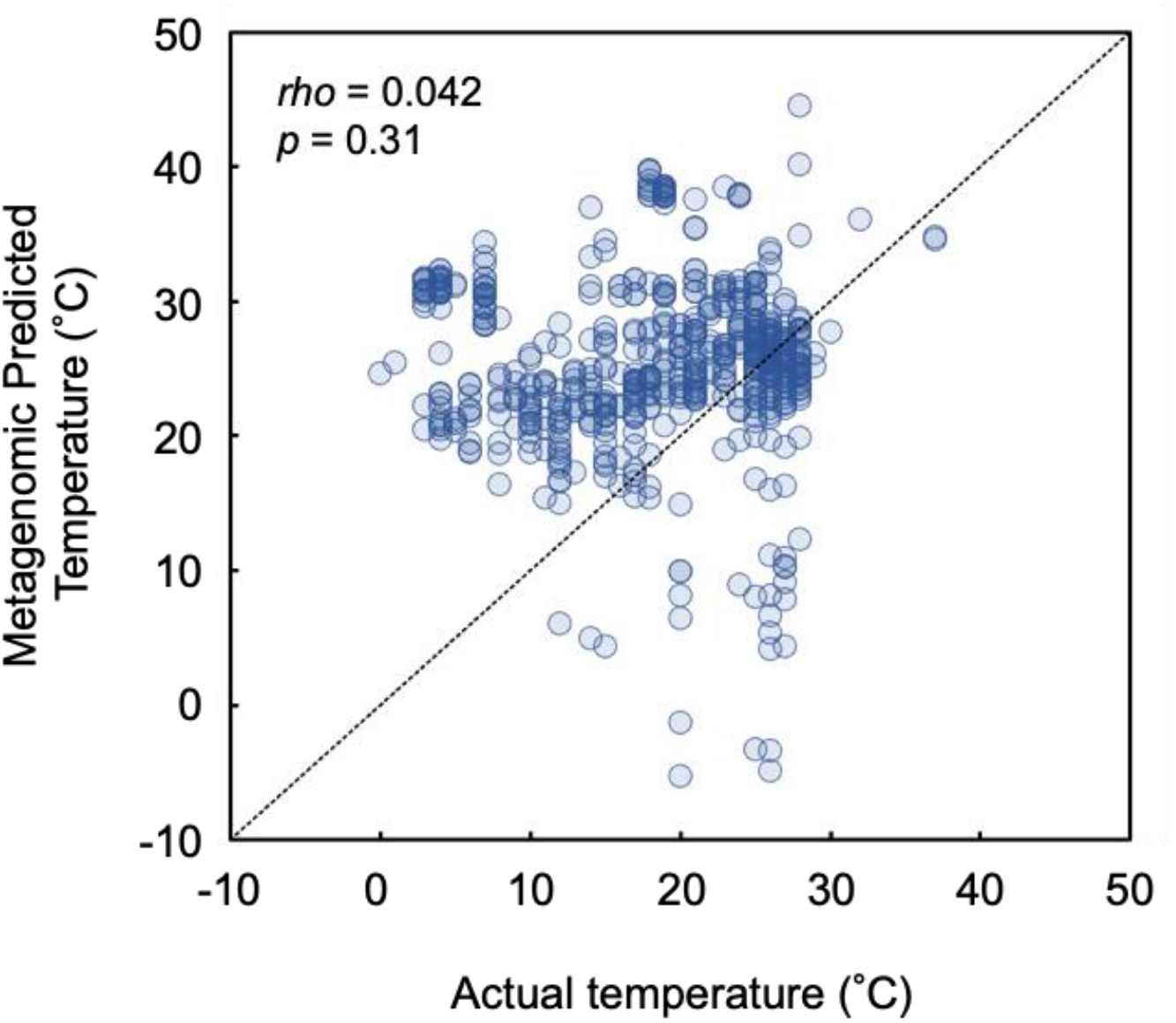
Temperature prediction based on data for the ocean metagenome. Points represent data for 586 metagenomic samples of ocean water. From data stored in the MicrobeDB database, the sequence data containing “ocean”, “sea”, or “deep sea” for Metagenome/Microbes Environmental Ontology was used. The dotted line represents the perfect match between the actual and predicted temperatures. The Spearman’s correlation coefficient and *p*-value are shown.

